# Therapeutic effects of favipiravir against severe fever with thrombocytopenia syndrome virus infection in a lethal mouse model: dose-efficacy studies upon oral administration

**DOI:** 10.1101/399899

**Authors:** Hideki Tani, Takashi Komeno, Aiko Fukuma, Shuetsu Fukushi, Satoshi Taniguchi, Masayuki Shimojima, Akihiko Uda, Shigeru Morikawa, Nozomi Nakajima, Yousuke Furuta, Masayuki Saijo

**Affiliations:** Department of Virology I, National Institute of Infectious Diseases, Tokyo, Japan; Department of Virology, Graduate School of Medicine and Pharmaceutical Sciences, University of Toyama, Toyama, Japan; Research Laboratories, Toyama Chemical Co., Ltd., Toyama, Japan; Department of Veterinary Science, National Institute of Infectious Diseases, Tokyo, Japan

## Abstract

**Background:** Severe fever with thrombocytopenia syndrome (SFTS), caused by SFTS virus (SFTSV), is a viral hemorrhagic fever with a high case fatality rate. Favipiravir was reported to be effective in the treatment of SFTSV infection *in vivo* in type I interferon receptor knockout (IFNAR^-/-^) mice at treatment dosages of both 60 mg/kg/day and 300 mg/kg/day for a duration of 5 days.

**Methods:** In this study, the efficacy of favipiravir at dosages of 120 mg/kg/day and 200 mg/kg/day against SFTSV infection in an IFNAR^-/-^ mouse infection model was investigated. IFNAR^-/-^ mice were subcutaneously infected with SFTSV at a 1.0 × 10^6^ 50% tissue culture infectious dose followed by twice daily administration of favipiravir, comprising a total dose of either 120 mg/kg/day or 200 mg/kg/day. The treatment was initiated either immediately post infection or at predesignated time points post infection.

**Results:** All mice treated with favipiravir at dosages of 120 mg/kg/day or 200 mg/kg/day survived when the treatment was initiated at no later than 4 days post infection. A decrease in body weight of mice was observed when the treatment was initiated at 3–4 days post infection. Furthermore, all control mice died. The body weight of mice did not decrease when treatment with favipiravir was initiated immediately post infection at dosages of 120 mg/kg/day and 200 mg/kg/day.

**Conclusions:** Similar to the literature-reported peritoneal administration of favipiravir at 300 mg/kg/day, the oral administration of favipiravir at dosages of 120 mg/kg/day and 200 mg/kg/day to IFNAR^-/-^ mice infected with SFTSV was effective.

**Author summary:** Severe fever with thrombocytopenia syndrome (SFTS), which is caused by SFTS virus (SFTSV), is a generalized infectious disease with a high case fatality rate. Currently, no effective therapeutics for SFTS is available; therefore, the development of effective antiviral drugs is needed. Favipiravir exhibits antiviral activity against various RNA viruses, including SFTSV. The present study demonstrated the efficacy of favipiravir in the treatment of SFTSV infection in a lethal mouse model, when the dose was set similar to that approved for anti-influenza drug in humans by the Ministry of Health, Labour and Welfare, Japan. The present study suggests that favipiravir is a promising drug for the treatment of SFTSV infection.

## Introduction

Severe fever with thrombocytopenia syndrome (SFTS) is caused by SFTS virus (SFTSV), belonging to the family *Phenuiviridae* (genus *Phlebovirus*). SFTS is a viral hemorrhagic fever with a high case fatality rate; it was first reported as a novel infectious disease in China [1, 2], followed by discovery in South Korea and Japan [3, 4]. It is characterized by marked reduction in platelet, white blood cell, and total blood cell counts in patients. Hemorrhagic symptoms, such as gingival oozing, bloody diarrhea, and hematuria, are commonly observed in patients with severe and fatal SFTS [3, 5, 6]. Because of the associated high mortality rate, it is critical to develop specific and effective therapy for SFTS. Unfortunately, no such treatment has been developed yet. The inhibitory effect of ribavirin on the replication of SFTSV has been elucidated *in vitro* as well as *in vivo* [7, 8]. Although ribavirin inhibited the replication of SFTSV *in vitro* in a dose-dependent manner, therapeutic effect *in vivo* was limited in comparison with that of favipiravir. Thus, an anti-SFTSV effect of ribavirin is limited or absent in the clinical setting [9, 10]. Favipiravir is an RNA-dependent RNA polymerase inhibitor and a potent broad-spectrum antiviral drug. It inhibits the replication of multiple families of RNA viruses *in vitro* and *in vivo* [11, 12]. Favipiravir is a therapeutic antiviral drug against influenza virus approved in Japan. However, during the 2014–2015 Ebola outbreak in West Africa, it was also considered as a candidate agent against Ebola virus infection [13, 14]. In addition, favipiravir was demonstrated to have antiviral effects against the newly discovered emerging viruses SFTSV and Heartland virus (HRTV) [15]. HRTV is an emerging tick-borne virus, which, similar to SFTSV, belongs to the genus *Phlebovirus* in the family *Phenuiviridae*. Patients infected with HRTV show similar symptoms as SFTS patients. The efficacy of favipiravir against HRTV infections was demonstrated in animal infection models using STAT2 knockout hamsters [15].

Reportedly, favipiravir is effective when administered even after symptoms appeared. The antiviral effects of favipiravir against SFTSV were confirmed in a mouse model as well as STAT2 knockout hamster model [16]. We have previously demonstrated the antiviral effects of favipiravir against SFTSV in a lethal mouse model using IFNAR^-/-^ mice. In the study, the highest dose of favipiravir used in mice experiments, at which side effects did not appear, was 300 mg/kg/day via intraperitoneal (i.p.) route. All mice treated with favipiravir at 300 mg/kg/day survived without showing any symptoms upon SFTSV infection. In the mouse model, all mice also survived when treated i.p. with favipiravir at 60 mg/kg/day. However, their body weight decreased by approximately 10% [8]. In the present study, the efficacy of favipiravir in the mouse lethal model was evaluated at dosages of 120 mg/kg/day via oral administration (p.o.) and 200 mg/kg/day p.o. The two doses of favipiravir were selected in clinical trials to evaluate the efficacy of favipiravir against influenza virus infections in humans.

Favipiravir dosages of 120 mg/kg/day p.o. and 200 mg/kg/day p.o. have been applied for approval in Japan, and the phase III study conducted in the USA. The aim of this study was to assess the efficacy of favipiravir at dosages of 120 mg/kg/day p.o. and 200 mg/kg/day p.o. in the treatment of SFTSV infection in the lethal mouse model using IFNAR^-/-^ mice.

## Materials and methods

### Ethics statement

All animal experiments were performed in biological safety level 3 (BSL-3) containment laboratories at the National Institute of Infectious Diseases (NIID) in Japan and adhered to NIID regulations and guidelines on animal experimentation. Protocols were approved by the Institutional Animal Care and Use Committee of the NIID (No. 215024).

### Cells, viruses, and antiviral compounds

Vero cells obtained from American Type Culture Collection (Summit Pharmaceuticals International, Japan) were maintained in Dulbecco’s modified Eagle’s medium (DMEM) supplemented with 10% heat-inactivated fetal bovine serum and antibiotics (DMEM-10FBS). The SFTSV Japanese strain SPL010 was used in this study [8]. Pseudotyped vesicular stomatitis viruses (VSV) possessing SFTSV-GP or VSV-G, designated SFTSVpv or VSVpv, respectively, were used [17]. SPL010 virus stocks were stored at −80°C until use. All work with SFTSV was performed in BSL-3 containment laboratories in the NIID in accordance with the institutional biosafety operating procedures. Favipiravir (Toyama Chemical Co., Ltd., Toyama, Japan) was suspended in 0.5% (*w*/*v*) methylcellulose solution.

### Animal experiments

IFNAR^-/-^ C57BL/6 mice were produced as described previously [8]. IFNAR^-/-^ C57BL/6 mice were bred and maintained in an environmentally controlled specific pathogen-free animal facility of the NIID. Eight- to 10-week-old male mice were used. Favipiravir was administered in mice using a stomach probe after subcutaneous inoculation (s.c.) with 1.0 × 10^6^ 50% tissue culture infectious dose (TCID_50_) of SFTSV in 100 μl DMEM. Treatments were commenced at 1 h post infection or at 1, 2, 3, 4, or 5 days post infection and continued for 5 days.

To determine the efficacy of favipiravir in the treatment of SFTSV infection, the mice were treated with favipiravir at dosages of either 120 mg/kg/day p.o. or 200 mg/kg/day p.o. [60 or 100 mg/kg/bis in die (BID), p.o.] for 5 days starting at various time points as described above (Fig.1). Blood samples (20 μl/animal) were obtained via tail vein puncture at intervals of 2–4 days over a period of 14 days (<4 blood drawings in total) for the measurement of viral RNA levels. Body weight was recorded daily for 2 weeks, and each mouse was monitored daily for the development of clinical symptoms such as hunched posture, ruffled fur, activity, response to stimuli, and neurological signs. When mice showed serious clinical symptoms or weight loss of more than 30 %, they were considered to be reached the humane endpoint so that they were anesthetized.

**Fig 1.**
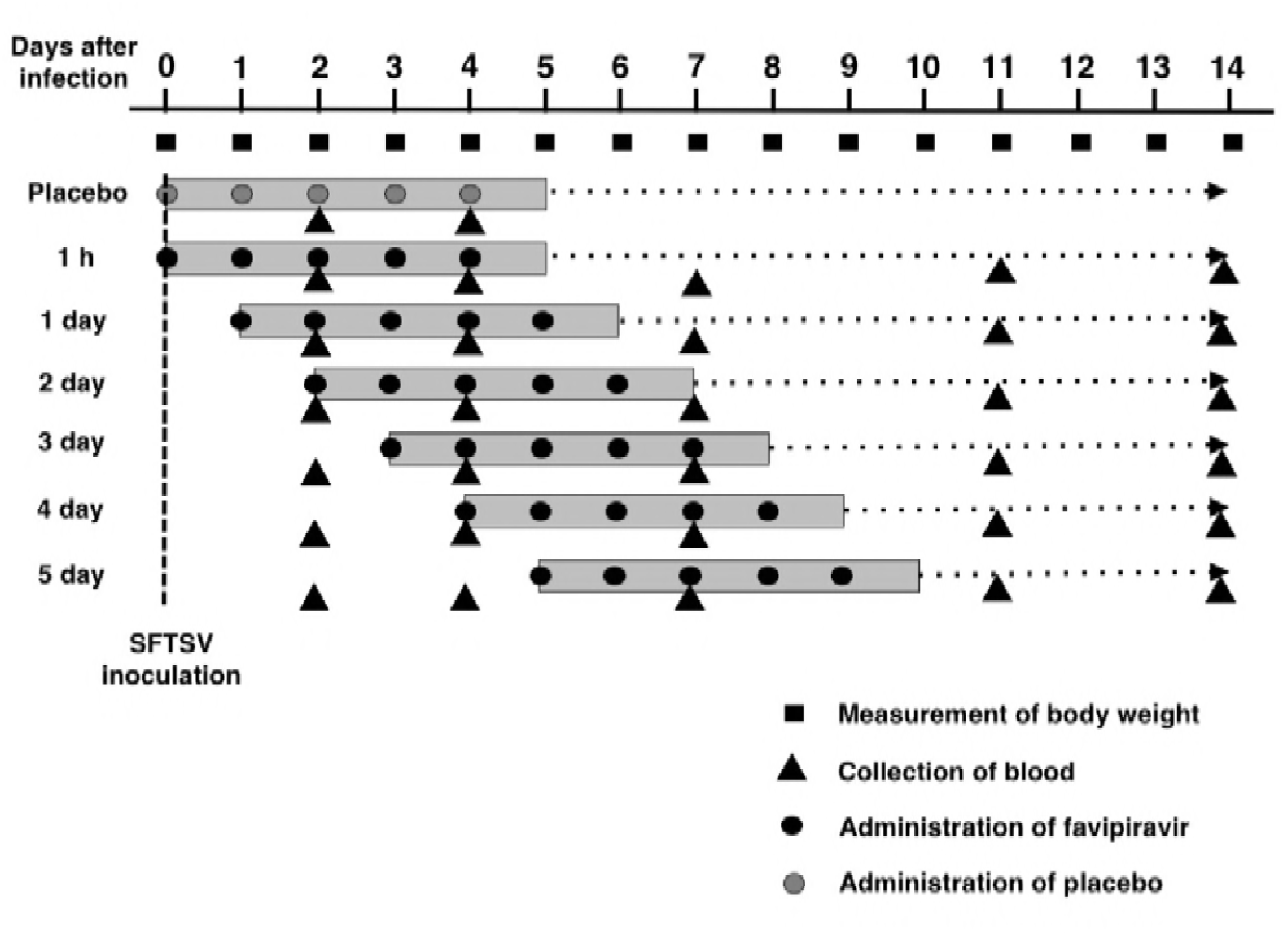
Schematic experimental design. Six mice in each group were administered favipiravir at either 120 mg/kg/day or 200 mg/kg/day starting at 1 h or 1, 2, 3, 4, or 5 days post infection and continued for 5 consecutive days. Placebo control mice were treated with an equal volume of 0.5% (w/v) methylcellulose solution administered at 1 h post infection and continued for 5 consecutive days.

### Viral RNA quantification

The concentration of SFTSV genomic RNA in blood was determined as previously described [18]. Total RNA was prepared from 20 μl of blood samples using High Pure Viral RNA Kit (Roche Diagnostics K.K., Tokyo, Japan). Gene expression was estimated using QuantiTect Probe RT-PCR kit (Qiagen, Hilden, Germany) according to the manufacturer’s protocol. Fluorescent signals were estimated using LightCycler 96 (Roche Diagnostics K.K., Tokyo, Japan). Statistical analyses were performed using GraphPad Prism6 Software. One-way analysis of variance (ANOVA) with Bonferroni’s multiple comparison test was used.

### Neutralization assay

The day of SFTSV infection was considered as Day 0 and days post infection were subsequently counted. Sera from the mice at a convalescent phase were obtained at Day 14. To examine the neutralization antibody responses against SFTSV of the mice at a convalescent-phase, pseudotyped VSV system was employed. SFTSVpv and VSVpv were pre-incubated with serially diluted sera of the mice at a convalescent-phase for 1 h at 37°C. Then, Vero cells were inoculated with each of the virus–serum mixtures. After 2 h of adsorption at 37°C, cells were washed with DMEM-10FBS and infectivity was determined by measuring luciferase activity after 24 h of incubation.

## Results

### Therapeutic efficacy of favipiravir against SFTSV infection in IFNAR^-/-^ mice

Consistent with the results of a previous study, the optimal lethal infectious dose of SFTSV strain SPL010 in mice was determined to be 1.0 × 10^6^ TCID_50_ [8]. All mice treated with favipiravir at dosages of 120 mg/kg/day or 200 mg/kg/day survived from a lethal SFTSV infection when treatment was initiated within 3 days and 4 days post infection, respectively (Fig. 2B and 2C). All control mice, infected with SFTSV died within 8 days post infection [8] (Fig. 2A). When treatment was initiated on Day 4, the mice treated with favipiravir at dosages of 120 mg/kg/day and 200 mg/kg/day exhibited 67% and 100% survival, respectively. However, under these conditions, the health of mice was highly deteriorated, with more than 15% weight loss. A few mice treated with favipiravir at a dosage of 200 mg/kg/day dose initiated on Day 5 survived even with 30% weight loss (Fig. 2C).

**Fig 2.**
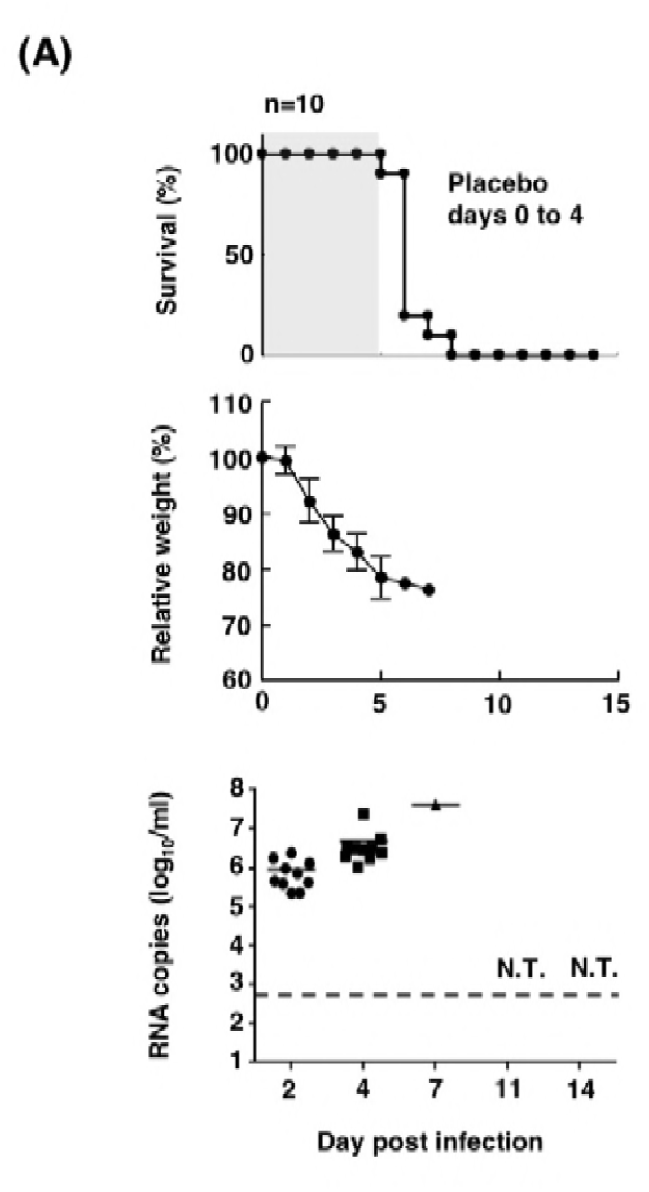
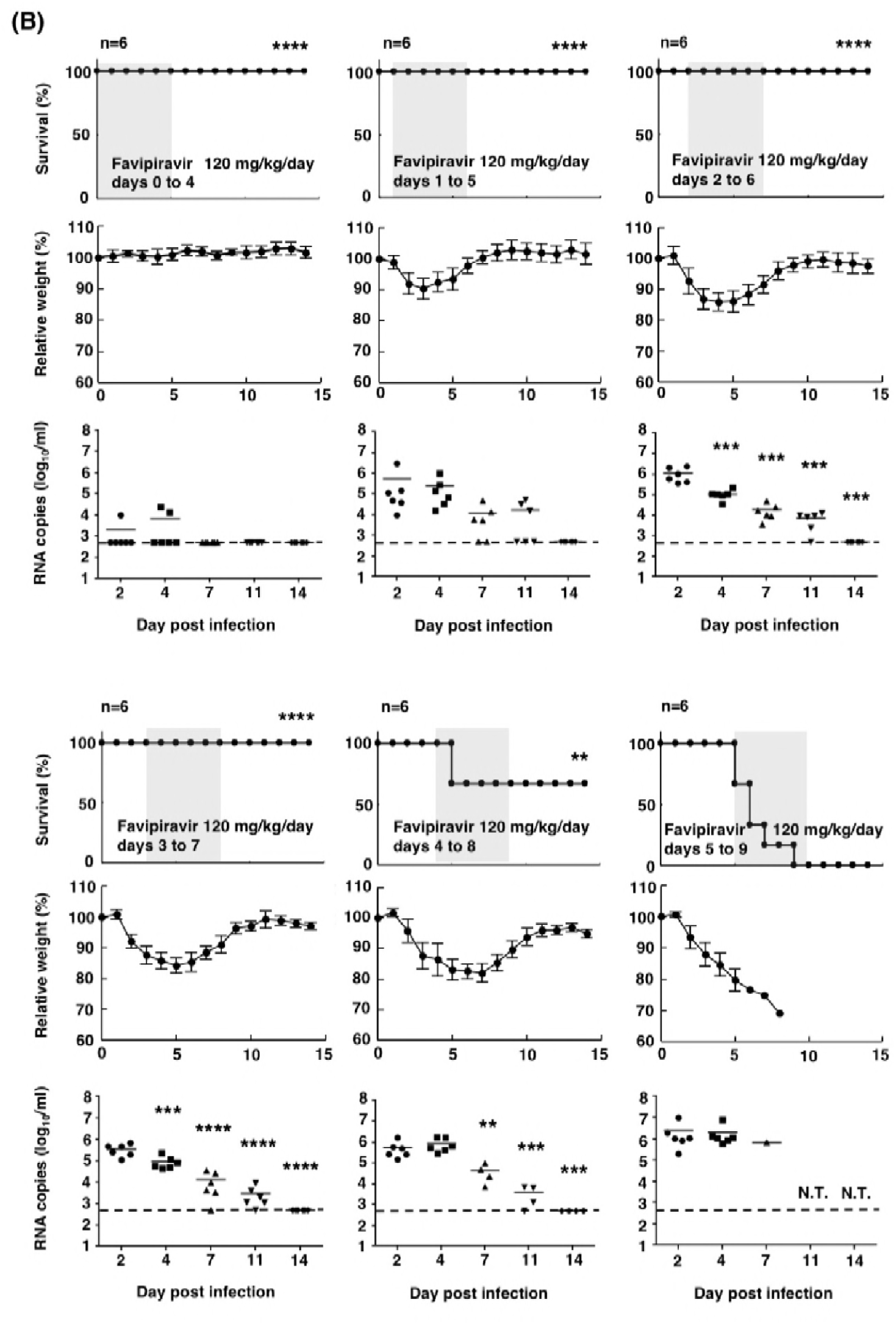
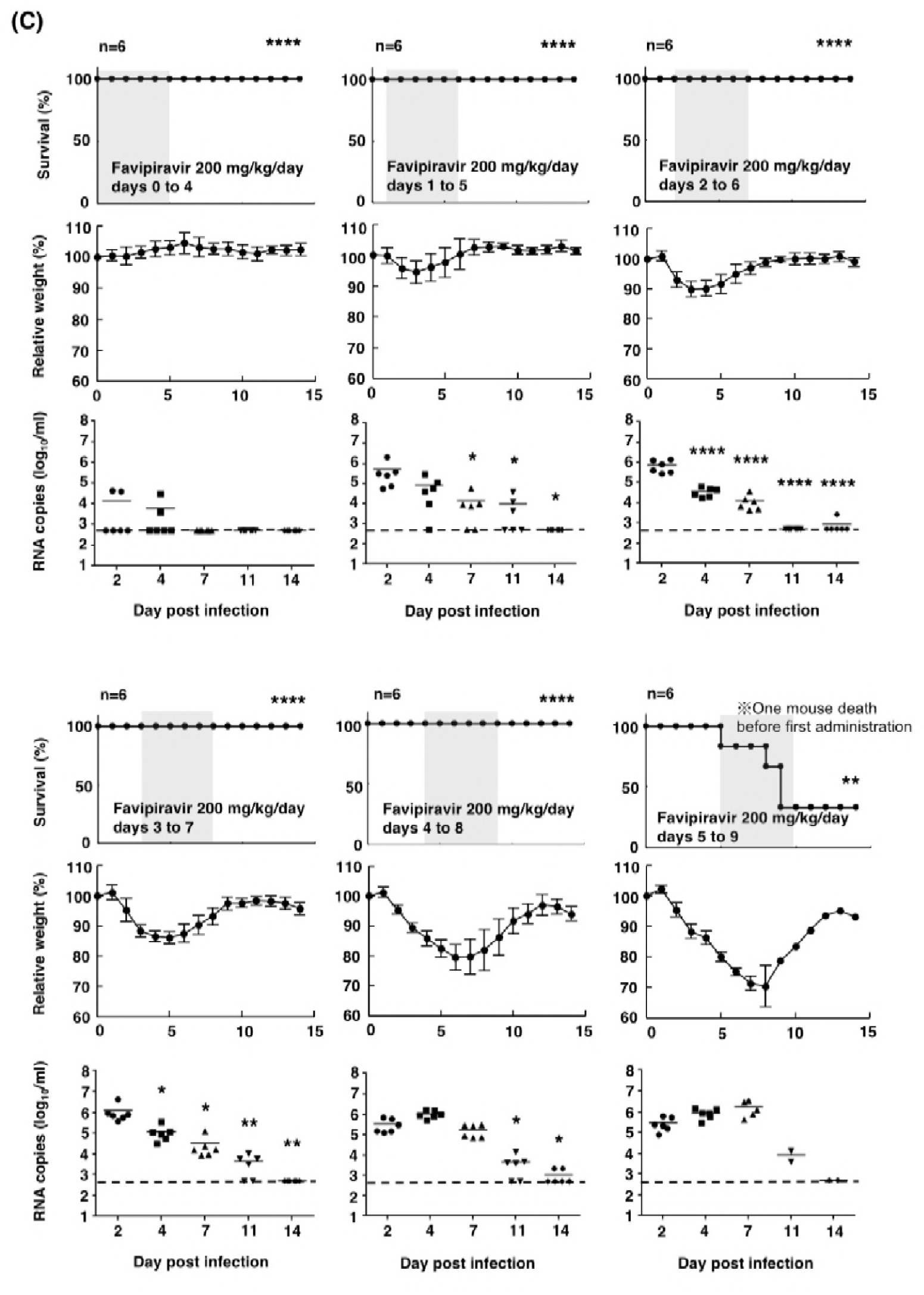
Effects of treatment with favipiravir against SFTSV infection in IFNAR^-/-^ mice. (A) Ten mice in the placebo control group were inoculated s.c. with 1.0 × 10^6^ TCID_50_ of SFTSV strain SPL010. Control mice received 0.5% (w/v) methylcellulose solution via the p.o. route. (B, C) Six mice in each group were inoculated s.c. with 1.0 × 10^6^ TCID_50_ of SFTSV strain SPL010. Mice were treated with favipiravir at a dose of 120 mg/kg/day (B, 60 mg/kg/BID, p.o.) or 200 mg/kg/day (C, 100 mg/kg/BID, p.o.). Treatment was commenced at 1 h or 1, 2, 3, 4, or 5 days post infection. Favipiravir was administered twice daily p.o. using a stomach probe until death or for 5 days as indicated in the upper columns (shaded in gray with survival curves). Survival was determined using Kaplan– Meier analysis and GraphPad Prism6 (GraphPad Software) and shown in the upper columns. Relative weights are shown as means with standard deviations (middle columns). SFTSV RNA levels in blood samples collected at 2, 4, 7, 11, or 14 days post infection were determined by quantitative RT-PCR assays (lower columns). One way ANOVA with Bonferroni’s multiple comparison test was used to determine statistical significance. Dashed lines indicate the detection limits of the assay in blood samples. Significance was determined in comparison to the results of the placebo group (for survivals) or Day 2 blood samples (for RNA copies): ****, *P* < 0.0001; ***, *P* < 0.001; **, *P* < 0.01; * *P* < 0.05; N.T., not tested.

The RNA levels in the blood of mice gradually decreased upon administration of favipiravir at dosages of 120 mg/kg/day and 200 mg/kg/day, respectively (Fig. 2B and 2C). There was no significant difference in the RNA levels between the two treatment groups. The viral RNA in blood was undetectable by Day 14 in most mice treated with favipiravir at dosages of 120 mg/kg/day and 200 mg/kg/day (Fig. 2B and 2C).

### Neutralizing antibody responses against SFTSV in the mouse sera at a convalescent-phase

To examine whether neutralizing antibodies were induced in the mice at a convalescent-phase, serum samples collected on Day 14 were tested for neutralizing activity with an assay using a pseudotyped VSV system. Sera of convalescent-phase mice neutralized SFTSVpv infection at a dilution of 1 in 800 (Fig. 3) and in a dilution-dependent manner (data not shown), whereas no significant neutralization of VSVpv infection was observed (Fig. 3). The induction of neutralizing antibody responses in mice wherein treatment was initiated on Days 0 or 5 seemed lower than the induction of neutralizing antibody responses in mice wherein treatment was initiated on Day 1 at a dosage of 200 mg/kg/day (Fig. 3B).

**Fig 3.**
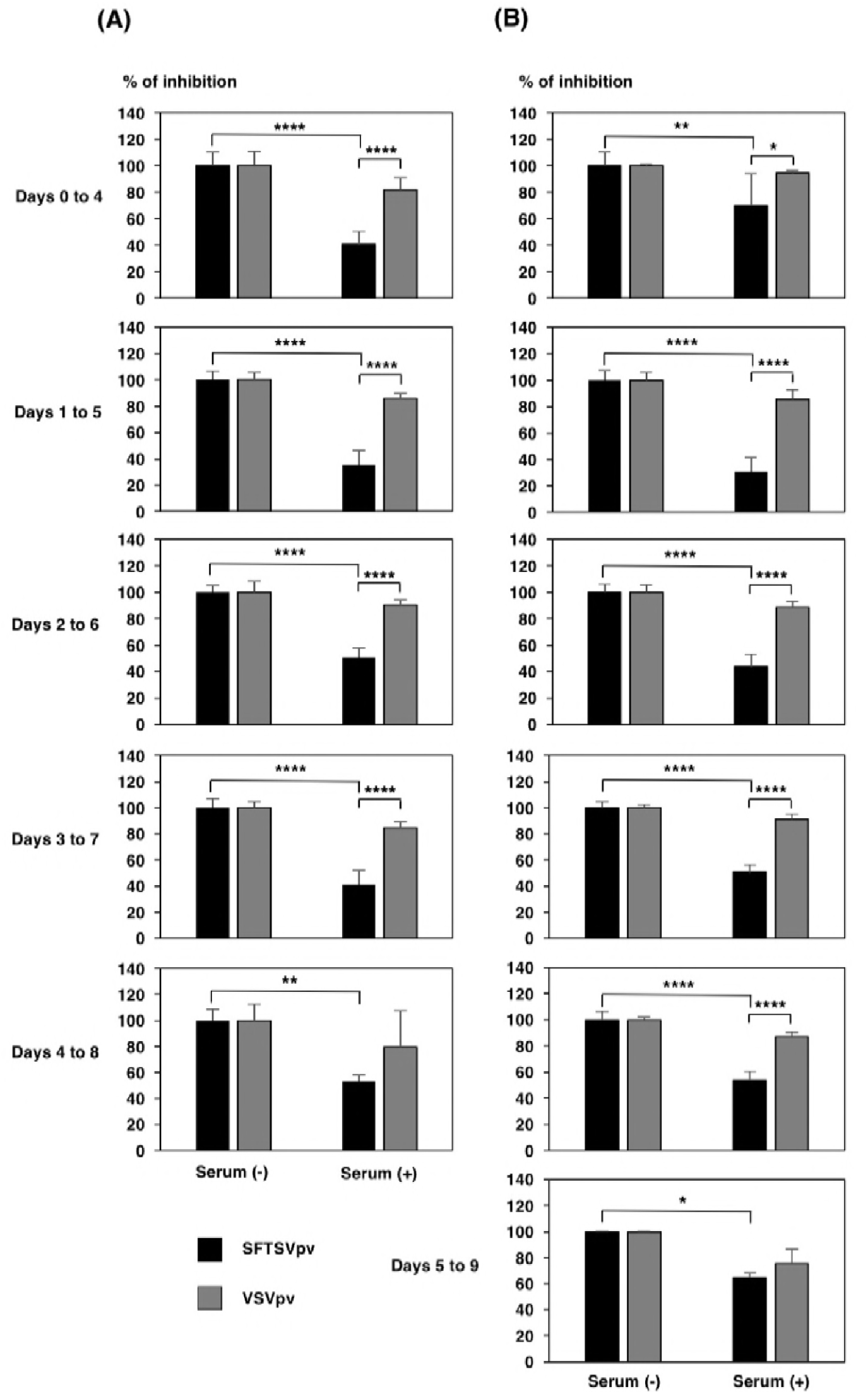
Neutralization of SFTSVpv by convalescent-stage mouse sera. SFTSVpv were preincubated with 800-fold diluted mouse sera collected on Day 14 (120 mg/kg/day treatment group [(A) left columns] and 200 mg/kg/day treatment group [(B) right columns]). Subsequently, Vero cells were infected with SFTSVpv. Infectivity of SFTSVpv was determined by measuring luciferase activities at 24 h post infection. Results from three independent assays are shown, with error bars representing standard deviations. Significance was determined in comparison to the results from non-serum treatment or infectivity of VSVpv. ****, *P* < 0.0001; **, *P* < 0.01; * *P* < 0.05.

## Discussion

We have previously demonstrated the protective efficacy of favipiravir in the treatment of SFTSV infection at dosages of 300 mg/kg/day i.p. in the lethal mouse model [8]. Since favipiravir is approved for anti-influenza drug as a formula of p.o. drug in Japan, we have tested the efficacy of favipiravir at dosages of 120 mg/kg/day and 200 mg/kg/day p.o. against SFTSV in the lethal mouse model. The results demonstrated favipiravir at both dosages were effective via oral administration. The dosages were the standard dose applicable in humans. For utilizing favipiravir as an anti-influenza drug in humans, a dosage of 120 mg/kg/day p.o. has been set for clinical use in Japan and a dosage of 200 mg/kg/day p.o. has been set for phase III studies in the USA, respectively. With regard to the Ebola virus disease (EVD) outbreak that occurred in West Africa in 2013–2015, favipiravir was required to be administered at a higher dose for the treatment of EVD than that required for the treatment of influenza. This was based on the higher IC_50_ values of favipiravir for Ebola virus *in vitro* and *in vivo* [14, 19, 20].

The effective concentration of favipiravir in blood is considered to be similar when administered p.o. and when administered i.p. [21]. Here the therapeutic effect of favipiravir in the treatment of SFTSV infection was observed both when administered p.o. as well as when administered i.p. In contrast to the previous reports, where favipiravir was administered once a day, favipiravir was administered twice a day (BID) in the present study. The antiviral effects of favipiravir when administered orally at the tested doses might be higher than those when administered via the intraperitoneal route quaque die [8]. This difference may be attributed to the maintenance of effective favipiravir concentration in blood. Furthermore, the observed therapeutic effect was obtained not only due to a direct inhibition of viral replication by favipiravir but also due to the production of neutralizing antibodies against SFTSV in the later phase of the disease (Fig. 3). The neutralizing antibody responses were higher in mice wherein treatment was initiated on Days 1 and 2 than in those wherein treatment was initiated on Day 0. This may be attributed to the amount of replicated virus as an antigen. Conversely, the production of neutralizing antibodies was weak in mice wherein treatment was initiated on Day 5, suggesting that neutralizing antibody producing cells were more heavily damaged in mice wherein the treatment was initiated in the later stages of the disease.

The therapeutic effect of favipiravir is remarkably higher against SFTS in animal models than other reported viral infectious diseases [19, 22, 23]. Administration of favipiravir after the onset of the disease did not show any efficacy in the treatment of EVD or Crimean-Congo hemorrhagic fever viral infection in animal models [19, 22, 23]. Conversely, the administration of favipiravir in the mice infected with SFTSV within 4 days post infection showed efficacy even at a dosage of 120 mg/kg/day, which is the dosage approved to be prescribed to humans (Fig. 2). Therefore, favipiravir was effective not only for prophylactic use but also for treating SFTS in the mouse model. However, it was too late to initiate the administration of favipiravir at Day 5 in the mice model (Fig. 2). The results obtained in the present study indicate that favipiravir should be administered as early as possible post infection. This also indicates that favipiravir should be administered as early as possible from disease onset for the treatment of patients with SFTS.

Currently, there is no antiviral therapy available for the treatment of SFTSV infection. Here, we studied the efficacy of favipiravir at dosages of 120 mg/kg/day p.o. and 200 mg/kg/day p.o. in the treatment of mice infected with SFTSV. These dosages can also be applied to humans. Currently, clinical trials are underway for evaluating the efficacy of favipiravir in the treatment of patients with SFTS in Japan [24]. We hope that favipiravir will not only be used as a prophylactic drug against SFTS in the near future but also as a therapeutic drug in clinical practice.

## Acknowledgments

We gratefully acknowledge Ms. Momoko Ogata, Ms. Junko Hirai, and Ms. Kaoru Hounoki for their technical and secretarial assistances.

